# Click-to-Release: Cleavable Radioimmunoimaging with ^89^Zr-DFO-*Trans*-Cyclooctene-Trastuzumab Increases Tumor-to-Blood Ratio

**DOI:** 10.1101/2023.03.27.534155

**Authors:** Maria Vlastara, Raffaella Rossin, Freek J.M. Hoeben, Kim E. de Roode, Milou Boswinkel, Laurens H.J. Kleijn, James Nagarajah, Mark Rijpkema, Marc S. Robillard

## Abstract

One of the main challenges of PET imaging with ^89^Zr-labeled monoclonal antibodies (mAbs) remains the long blood circulation of the radiolabeled mAbs, leading to high background signals, decreasing image quality. To overcome this limitation, here we report the use of a bioorthogonal linker cleavage approach (click-to-release chemistry) to selectively liberate [^89^Zr]Zr-DFO from *trans*-cyclooctene-functionalized trastuzumab (TCO-Tmab) in blood, following the administration of a tetrazine compound (trigger) in BT-474 tumor-bearing mice.

**Methods:** We created a series of TCO-DFO constructs and evaluated their performance in [^89^Zr]Zr-DFO release from Tmab in vitro using different trigger compounds. The in vivo behavior of the best performing [^89^Zr]Zr-TCO-Tmab was studied in healthy mice first, to determine the optimal dose of the trigger. To find the optimal time for the trigger administration, the rate of [^89^Zr]Zr-TCO-Tmab internalization was studied in BT-474 cancer cells. Finally, the trigger was administered 6 h or 24 h after [^89^Zr]Zr-TCO-Tmab-administration in tumor-bearing mice to liberate the [^89^Zr]Zr-DFO fragment. PET scans were obtained of tumor-bearing mice that received the trigger 6 h post-[^89^Zr]Zr-TCO-Tmab administration.

**Results:** The [^89^Zr]Zr-TCO-Tmab and trigger pair with the best in vivo properties exhibited 83% release in 50 % mouse plasma. In tumor-bearing mice the tumor-blood ratios were markedly increased from 1.0 ± 0.4 to 2.3 ± 0.6 (p=0.0057) and from 2.5 ± 0.7 to 6.6 ± 0.9 (p<0.0001) when the trigger was administered at 6 h and 24 h post-mAb, respectively. Same day PET imaging clearly showed uptake in the tumor combined with a strongly reduced background due to the fast clearance of the released [^89^Zr]Zr-DFO-containing fragment from the circulation through the kidneys.

**Conclusions:** This is the first demonstration of the use of *trans*-cyclooctene-tetrazine click-to-release chemistry to release a radioactive chelator from a mAb in mice to increase tumor-blood ratios. Our results suggest that click-cleavable radioimmunoimaging may allow for substantially shorter intervals in PET imaging with full mAbs, reducing radiation doses and potentially even enabling same day imaging.

## Introduction

Monoclonal antibodies (mAbs) are great targeting agents with high uptake in tumors as a consequence of their long blood circulation [1]. A variety of mAb-based biopharmaceuticals are used in cancer imaging and therapy [2]. For cancer imaging using positron emission tomography (PET) and long-circulating mAbs, the radionuclide of choice is β^+^ emitting Zr-89, with its 3.3 days half-life [3,4]. Some limitations to using ^89^Zr-labeled mAbs include i) the high radiation dose in excretory organs such as the liver [5,6] and ii) the need for a 4-to-7-day interval between injection of radioactivity and the PET scan to achieve sufficiently high tumor-to-blood ratios for imaging [7,8].

A few solutions have been proposed to address these issues. Short circulating mAb fragments have been investigated, but they can exhibit suboptimal pharmacokinetics with a decreased tumor uptake due to fast renal clearance, and high kidney retention [9]. Another approach is tumor pretargeting, wherein injection of a tagged mAb is followed by a small radiolabeled probe that binds the tag in the tumor or otherwise clears rapidly from blood [10]. However, this method is mainly limited to the targeting of non-internalizing receptors and it typically requires the use of a clearing agent to remove the tagged mAb from circulation to the liver before probe administration, increasing the complexity of the method [11,12]. While clearing agents have also been used to remove radiolabeled mAbs from blood once the desired tumor uptake has been achieved, this approach results in high radioactivity retention in liver [13–15].

It would be advantageous to be able to separate the radioactive moiety from the mAb in blood and other non-target tissues, after sufficient tumor uptake has occurred, allowing rapid renal clearance of the radioactivity, boosting tumor-to-background ratios. mAb-chelate conjugates cleavable by endogenous or exogenous enzymes have been developed [16-18]. However, the former approach was designed to reduce the radioactivity in liver instead of in blood [16, 17] and the latter approach was hampered by the need for multiple doses of enzyme with the risk of immunogenicity [18]. We set out to develop a method where the separation of the radioactivity from the mAb occurs in blood and is induced by an external stimulus based on a bioorthogonal reaction, enabling a temporal and potentially spatial control over the cleavage (Figures 1 and 2).

**Figure 1.**
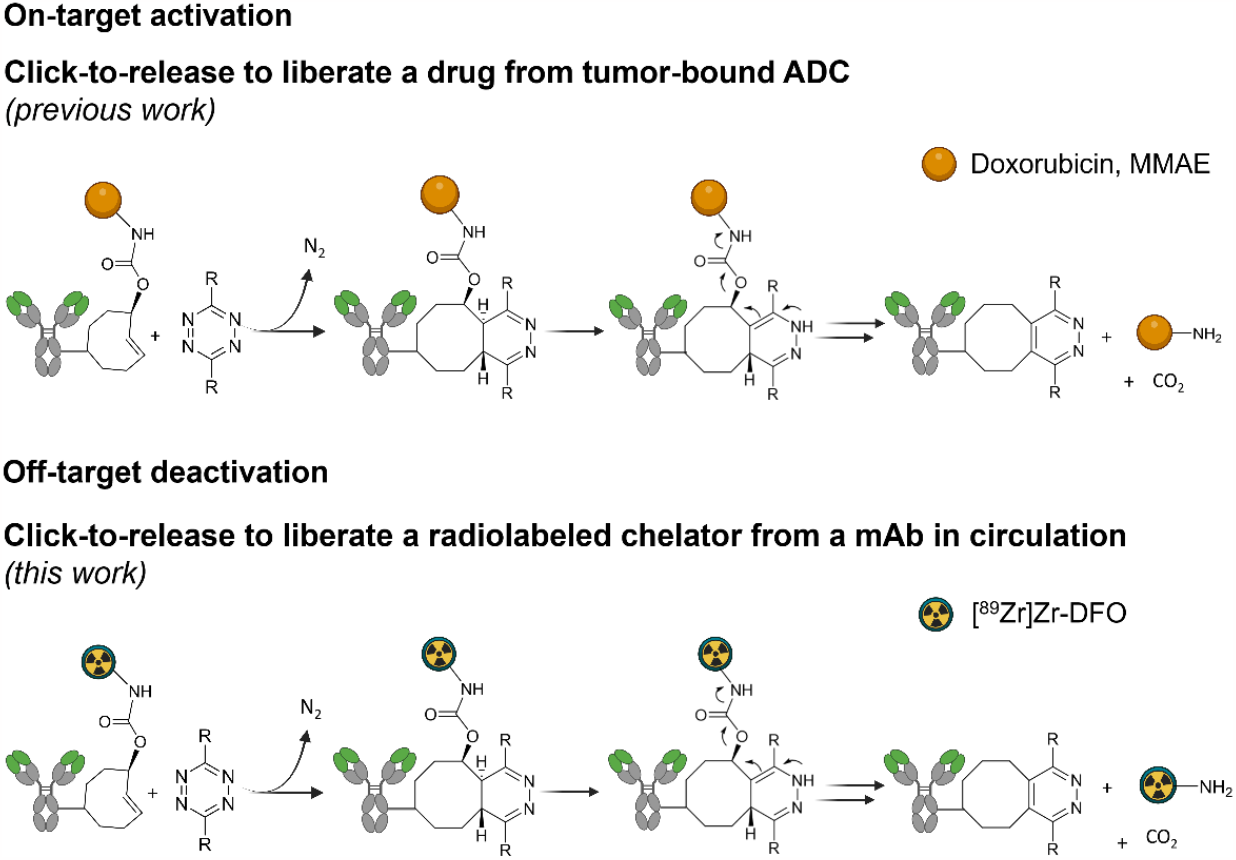
Click-to-release concept. Top: inverse electron-demand Diels-Alder (IEDDA) pyridazine elimination reaction between TCO and tetrazine, wherein the released payload is doxorubicin or MMAE (previous work) [19]; bottom: IEDDA pyridazine elimination reaction wherein the released payload is a radioactive moiety (this work).

**Figure 2.**
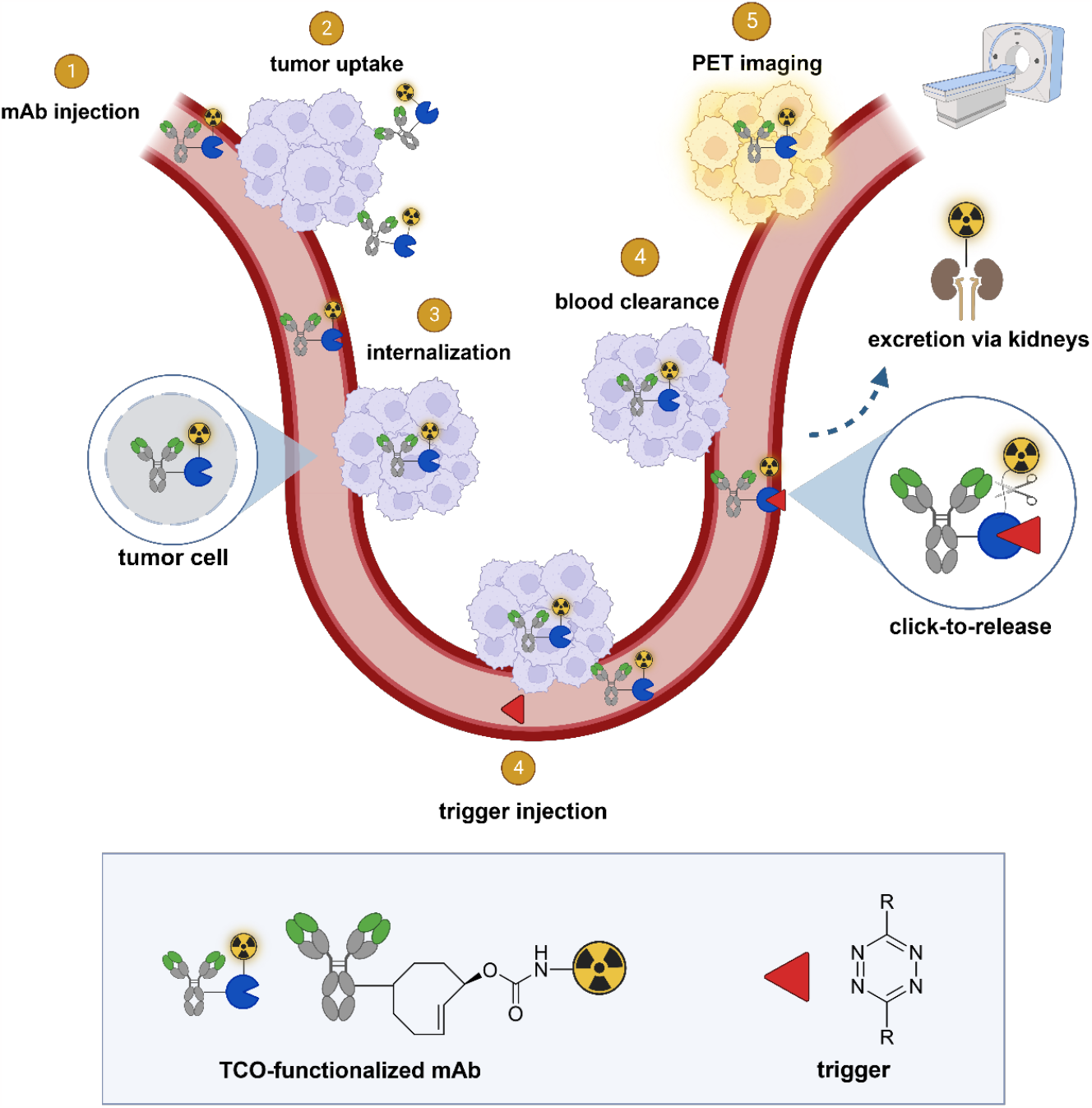
General concept of cleavable radioimmunoimaging using click-to-release chemistry in vivo. The TCO-linked and radiolabeled mAb is administered and internalizes in the tumor cells. Once enough tumor internalization has occurred, the trigger (tetrazine) is administered, which can solely react with TCO in blood and in other extracellular compartments given its non-cell-permeable nature. The result of this bioorthogonal reaction is the rapid release of the radiolabeled chelator in circulation and its efficient renal excretion, increasing the tumor-blood (T/B) ratio. Created with BioRender.com

We previously developed a bioorthogonal cleavage reaction (Figure 1) based on the inverse electron-demand Diels-Alder (IEDDA) reaction between a *trans*-cyclooctene (TCO) and a tetrazine [19]. This IEDDA pyridazine elimination was used for on-target activation of non-internalizing chemically-cleavable antibody-drug conjugates (ADCs), wherein a TCO linker releases the drug from the allylic position upon reaction with a separately administered tetrazine in the extracellular tumor microenvironment (Figure 1) [20,21]. Based on the high and selective in vivo reactivity combined with good stability that we observed for the chemically-cleavable ADC, we hypothesized that this reaction could be equally effective for controlled off-target deactivation and started to develop this in the context of click-cleavable radioimmunoimaging and - therapy [22].

In this approach, a mAb targeting an internalizing receptor is conjugated with a TCO-linked chelator, radiolabeled, and administered intravenously (i.v.). After sufficient tumor accumulation and internalization has occurred, a non-cell permeable tetrazine, referred to as “trigger”, is administered in a second step. The trigger reacts solely with the mAb in blood and in other extracellular domains, resulting in the release of the radiolabeled chelator, which as a small fragment clears rapidly via the kidneys while the intracellular radioactivity in the tumor is retained (Figure 2). Zhang and co-workers recently reported a similar approach using in vivo NO generation from the vasodilating drug glyceryl trinitrate (GTN) as the trigger for the release of the radionuclide ^131^I from a nanoparticle [23]. While elegant, it is not clear whether the pharmacodynamics of the drug itself limits its applicability as a trigger molecule for the present application, and whether the NO levels are sufficiently high in plasma (i.e. in the extracellular blood compartment).

Here we describe the development of click-cleavable radiolabeled trastuzumab (Tmab), which binds the internalizing HER2 receptor [24,25]. ^89^Zr-labeled Tmab is used extensively to image HER2-positive cancer patients with PET [8,26,27]. We synthesized a small series of TCO-deferoxamine (TCO-DFO) linker-chelate constructs comprising a conjugation moiety and PEG spacers on one or both sides of the TCO to increase hydrophilicity and to potentially modulate the performance of the click-to-release reaction (Figure 3). Following conjugation to Tmab and ^89^Zr-labeling, trigger-induced [^89^Zr]Zr-DFO release was tested in vitro and the most promising [^89^Zr]Zr-TCO-Tmab/trigger pair was then evaluated in vivo, confirming that PET imaging of cancer with full mAbs can be significantly improved using click-cleavable radioimmunoconjugates.

**Figure 3.**
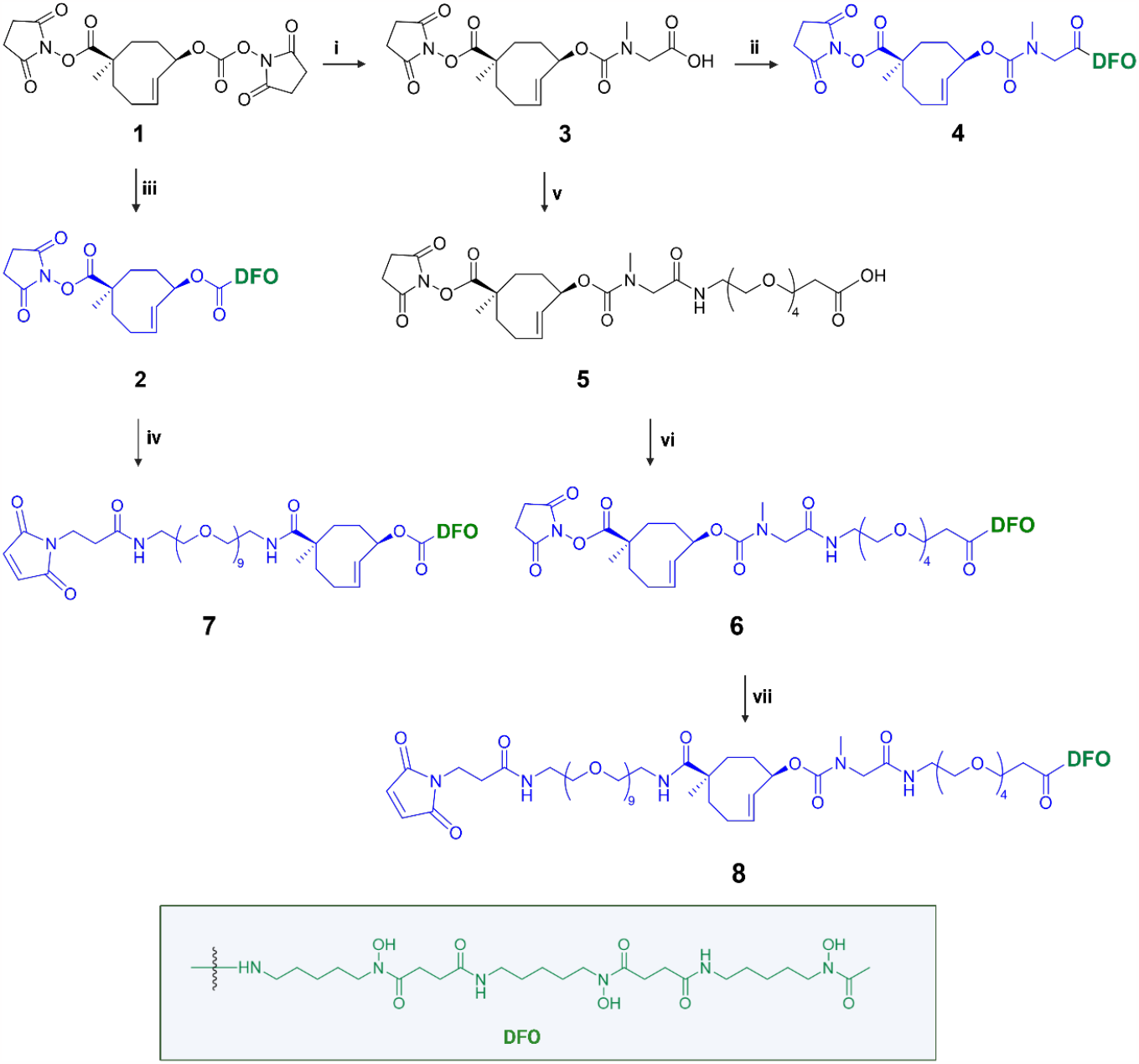
Synthesis of chemically-cleavable linker-chelates comprising a TCO functionalized on the releasing end (allylic position) with the [^89^Zr]Zr-DFO chelator. Reagents and conditions: (i) sarcosine, water, RT; (ii) PyBOP, DIPEA, deferoxamine mesylate salt, DMSO, RT; (iii) deferoxamine mesylate salt, DMSO, RT; (iv) mal-amido-PEG_9_-amine, DMF, DIPEA, RT; (v) PyBOP, DIPEA, DMF, amino-PEG_4_-acid RT; (vi) PyPOB, DIPEA, deferoxamine mesylate salt, DMSO, RT; (vii) mal-amido-PEG_9_-amine, DMF, DIPEA, RT.

## Materials and Methods

Additional methods, materials, corresponding figures, and tables are provided in the Supplementary Information, including synthesis and characterization of all new compounds, cell internalization experiments, in vitro release experiments, in vivo experiments and PET imaging.

### Antibody conjugations

Tmab in chelex-treated PBS was reacted with 35 equivalents of linker-chelate constructs **2, 4**, and **6** in PBS (pH adjusted to 8.5), to afford conjugates **Tmab-2, Tmab-4** and **Tmab-6**, respectively. Constructs **7** and **8** were conjugated to Tmab after partial mAb reduction with TCEP (2.3 eq) followed by incubation with 10 equivalents of the constructs in PBS pH 6.8 (in the presence of 2 mM EDTA) to afford the conjugates **Tmab-7** and **Tmab-8**, respectively. For all conjugates the excess of unreacted linker-chelate was removed by Size Exclusion Chromatography (SEC), with the UV lamp off, in order to obtain the final conjugates with >95% purity, as confirmed by analytical SEC. The amount of TCO per Tmab in the conjugates was measured using a tetrazine titration [21] and was found to range from 1.2 to 2.5. All the conjugate solutions were stored in aliquots at -80 °C.

### ^89^Zr labeling

The radiolabeling of all conjugates was performed following a published procedure with slight modifications [28]. ^89^Zr oxalate solution was combined with the Tmab conjugate solution (40 μg in 2 mg/mL) in a final volume of 200 μL with a pH between 7.2 – 7.4. The reaction vial was heated to 37 ^°^C in the dark for 30 min. The radiolabeling yield determined by iTLC varied between 80-95%. The radiolabeling mixture was purified using Zeba spin desalting columns (0.5 mL, 40 kDa MW-cut off). The purity of the recovered radiolabeled conjugates was >98% as determined by iTLC, SDS-PAGE and FPLC.

### Release experiments in vitro

The release experiments in PBS were carried out in triplicate mixing 1.6 μg of [^89^Zr]Zr-TCO-Tmab (0.13 MBq) with an excess of trigger (300 eq, from 1 mM solution in DMSO). Before analyzing the reaction mixture using SEC, 50 μL of a 0.6% BSA solution in PBS was added. The release experiments in plasma were performed by mixing 1.6 μg of [^89^Zr]Zr-TCO-Tmab (0.13 MBq) with an excess of each trigger (300 eq, from 1 mM solution in DMSO), while the final volume was 1:1 (plasma: PBS).

### Animal studies

For animal experiments, the guidelines set by the Nijmegen and European Animal Experiments Committee were followed and all experiments were approved by the institutional Animal Welfare Committee of the Radboud University Nijmegen. Female BALB/c nude mice (7–9-week-old, 18–22 g body weight) were used. All animals were allowed to acclimate for 1 week before the start of the experiments. Upon arrival, the mice were identified with tattoos. All experiments were blinded.

For the experiments in tumor-bearing mice, on day 0 a pellet of 17β-estradiol (0.18 mg/pellet, 60-day release) was subcutaneously (s.c.) implanted in the mice followed by injection with BT-474 cells (5 million cells / mouse in 100 μL matrigel / RPMI 1:1) in the mammary fat pad, under anesthesia. Tumor size was determined by caliper measurements in three dimensions (tumor volume= 1/2 × 1 × w × h) twice per week and the tumors were allowed to grow for 3-4 weeks until the volume reached 30 - 50 mm^3^, at which point the animals were randomly allocated to the various groups. At euthanasia, blood was obtained by cardiac puncture and organs and tissues of interest were harvested, blotted dry, and weighed. The sample radioactivity was measured in a gamma counter (Wizard 1480, PerkinElmer) along with standards to determine the % injected dose per gram (% ID/g) and the % injected dose per organ (% ID/organ). Stomachs, small and large intestines were not emptied before γ-counting.

### Tmab conjugate blood kinetics in mice

Tumor-free mice (n=4) were injected intravenously (i.v.) with [^**89**^**Zr]Zr-Tmab-8** (0.5 mg/kg, ca 0.5 MBq in 100 μL saline per mouse). Blood was withdrawn via the vena saphena (ca 20 μL sample) at various time points between 1 h to 72 h post mAb injection.

Four days post mAb injection the mice were euthanize, one last blood sample was obtained via a cardiac puncture and selected organs and tissues were harvested for γ-counting. The half-lives in blood were calculated by fitting the data to bi-exponential curves (GraphPad Prism v 9).

### Tmab conjugate stability in mice

Tumor-free nude mice (n = 4) were injected i.v. with [^**89**^**Zr]Zr-Tmab-8** (0.5 mg/kg, ca 0.5 MBq in 100 μL saline per mouse). At selected time points (1 h, 3 h, 6 h 24 h, 48 h, 72 h and 96 h) blood samples were withdrawn from the vena saphena and collected in vials containing heparin. After radioactivity measurement, the blood samples were diluted with 200 μL PBS and the blood cells were removed by centrifugation. The supernatant was divided in two aliquots. One aliquot of supernatant was analyzed by SEC. To the other aliquot an excess of trigger was added and the solution was incubated at 37 ºC overnight followed by SEC analysis to quantify the extent of chelate release. The % release was normalized to the value obtained at t=0 and was plotted with time. The data were then analyzed using linear regression (GraphPad Prism, v. 9).

### Triggered Tmab conjugate cleavage in tumor-free mice

Tumor-free mice (n = 4) were injected i.v. with [^89^Zr]Zr-TCO-Tmab (0.5 mg/kg, ca 0.5 MBq in 100 μL saline per mouse), followed 1 h later by an i.v. dose of trigger (33.4 μmol/kg in 100 μL PBS with 5% DMSO). In the case of [^**89**^**Zr]Zr-Tmab-8**, one group of mice received one extra dose of trigger 2 h after the first trigger injection. Blood was withdrawn via the vena saphena (ca 20 μL sample) at various time points. The blood samples were weighted and were measured in a γ-counter along with standards. Mice were euthanized 24 h post-mAb injection and selected organs were harvested, weighted, and measured in the γ-counter.

### Triggered Tmab conjugate cleavage in tumor-bearing mice

Mice bearing BT-474 cancer xenografts (n = 5) were injected i.v. with [^**89**^**Zr]Zr-Tmab-8** (0.5 mg/kg, ca 0.5 MBq in 100 μL saline per mouse), followed at 6 h or 24 h post-mAb injection with an i.v. dose of trigger **10** (33.4 μmol/kg, in 100 μL PBS with 5% DMSO). Mice were euthanized 4 h after the trigger dose in both cases. The organs were harvested, weighted, and measured in the γ-counter.

### PET studies

Mice bearing BT-474 cancer xenografts (n = 4) were injected i.v. with [^**89**^**Zr]Zr-Tmab-8** (0.5 mg/kg, ca 5 MBq in 100 μL saline per mouse). After 5 h mice were imaged for 15 min under anaesthesia (2-3% isoflurane in air). One hour after recovery from anaesthesia the mice received a dose **10** i.v. (33.4 μmol/kg in 100 μL PBS with 5% DMSO) and, 4 hours later, the mice were anaesthetized again and imaged for 30 min. Inveon Acquisition Workspace software (version 1.5, Siemens Preclinical Solution, Erlangen, Germany) was used for PET scans reconstruction with an algorithm with shifted Poisson distribution and the following settings: matrix 256 × 256 × 161, pixel size 0.4 × 0.4 × 0.8 mm, with a corresponding beta of 0.05 mm.

### Data analysis

All data are presented as the mean ± standard deviation (SD). Curve fitting, linear regressions and area-under-the-curve calculations were performed with GraphPad Prism (v 9). Statistical analysis was performed using the Student’s t-test (two-tailed) with Welch’s correction and unpaired t-test using GraphPad Prism (v 9). Statistical significance was set at p < 0.05.

## Results and Discussion

### Design and synthesis of TCO-DFO linker-chelate conjugates

Detailed information for the synthesis of the TCO linker-chelate constructs is provided in the Supplemental Information. In this study we synthesized a series of chemically-cleavable linker-chelate conjugates (compounds **2, 4, 6, 7** and **8**) containing a TCO functionalized on the releasing end (allylic position) with DFO for ^89^Zr-labeling (Figure 3). All compounds were obtained starting from the bis-NHS-functionalized TCO (**1**) that was previously used for the development of chemically-cleavable ADCs [20, 21]. In linkers **6-8**, PEG spacers were introduced before the TCO, or between the TCO and the DFO chelator, or both (Figure 3) to increase the hydrophilicity of the linkers and to potentially influence the efficiency of the click-to-release reaction with the trigger. The N-methyl group on the carbamate in **4, 6, 7** and **8** was added to prevent a potential intramolecular side-reaction, which was found by Carlson *et al*. to produce a non-releasing species [29].

All conjugates were conjugated to Tmab either via NHS chemistry or maleimide chemistry producing **Tmab-2, Tmab-4, Tmab-6, Tmab-7** and **Tmab-8**, which were radiolabeled with Zr-89 to afford [^**89**^**Zr]Zr-Tmab-2, [**^**89**^**Zr]Zr-Tmab-4, [**^**89**^**Zr]Zr-Tmab-6, [**^**89**^**Zr]Zr-Tmab-7** and [^**89**^**Zr]Zr-Tmab-8**, respectively.

### Release experiments in vitro and in circulation in mice

The IEDDA pyridazine elimination reaction requires the use of tetrazines that upon reaction with the TCO linker efficiently afford the releasing 1,4-dihydropyridazine intermediate from the initially formed 4,5-dihydropyridazine cycloaddition product [19]. The highly reactive and biocompatible 3,6-bispyridyl-1,2,4,5-tetrazine motif, the tetrazine of choice for many in vivo applications such as pretargeted radioimmunotherapy [30], unfortunately does not afford an efficiently releasing 1,4-dihydropyridazine intermediate [19]. The 3,6-bismethyl-1,2,4,5-tetrazine (**9**, Figure 4) gives high release of TCO-conjugated payloads in vitro [19], however, its low reactivity and fast pharmacokinetics makes it less suitable for in vivo applications [21]. To harness the high reactivity of 3,6-bispyridyl tetrazine for improved in vivo efficacy of click-to-release approaches, we recently developed tetrazines with ortho-functionalized pyridyl substituents, such as the amide-substituted **10** (Figure 4), designed to promote the formation of the releasing 1,4-dihydropyridazine tautomer [31]. In contrast with the ca. 10% release achieved with the parent bispyridine-tetrazine [19], this compound yielded 80% in vitro antibody-drug conjugate (ADC) cleavage within 2 h increasing to near quantitative overnight, and exhibited efficient in vivo ADC cleavage as well.

**Figure 4.**
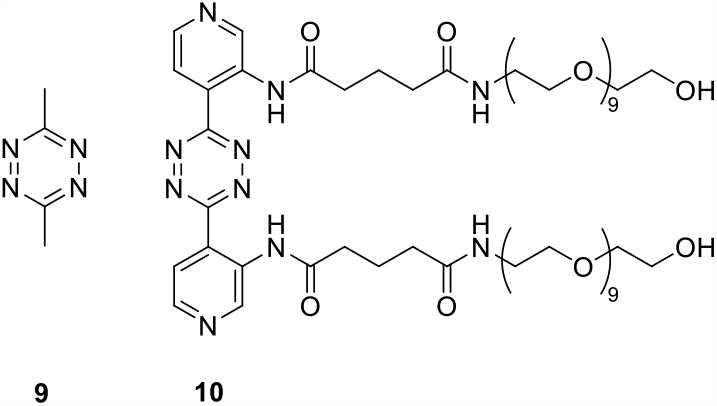
Triggers used in this study.

Compound **10** was evaluated as trigger for the cleavage of the [^89^Zr]Zr-TCO-Tmab conjugates in PBS for 24 h and compared with reference compound **9** (Table 1, Figure S1). While in general the release achieved with **10** was somewhat lower and the linker variations did not lead to significant differences, the highest release found with [^**89**^**Zr]Zr-Tmab-8** with 78% in PBS increased to 83% in 50% mouse plasma (Figure S2 and S3), which is close to what is maximally achievable with **9** [19-21]. Therefore, the combination of **10** with [^**89**^**Zr]Zr-Tmab-8** was selected for in vivo evaluation (Figure 5).

**Table 1.** Triggered release of radioactive fragment from [^89^Zr]Zr-TCO-Tmab conjugates in PBS. ^89^Zr-conjugates were mixed in PBS with an excess of trigger (300 eq) and incubated at 37 °C for 24 h. Samples were analyzed using SEC. The data represent the mean % ± SD (n=3).

| <sup>89</sup>Zr-Tmab | Release induced by triggers (mean% ± SD) |  |
| --- | --- | --- |
|  | **9** | **10** |
| [<sup>89</sup>Zr]Zr-Tmab-**2** | 80.4 ± 0.6 | 69.3 ± 1.7 |
| [<sup>89</sup>Zr]Zr-Tmab-**4** | 88.5 ± 0.3 | 75.1 ± 0.8 |
| [<sup>89</sup>Zr]Zr-Tmab-**6** | 86.9 ± 0.1 | 68.2 ± 4.2 |
| [<sup>89</sup>Zr]Zr-Tmab-**7** | 63.2 ± 2.3 | 59.4 ± 9.8 |
| [<sup>89</sup>Zr]Zr-Tmab-**8** | 89.5 ± 0.5 | 78.1 ± 0.6 |

**Figure 5.**
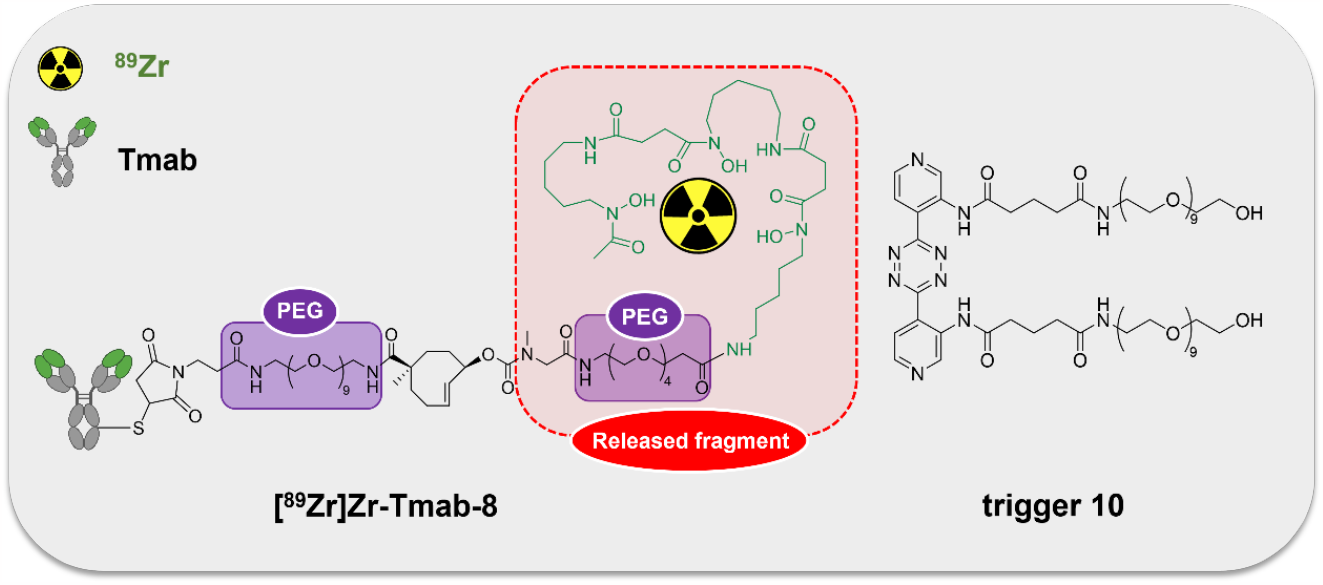
The ^89^Zr-TCO-Tmab and trigger pair used in the in vivo experiments. Chemical structure of [^**89**^**Zr]Zr-Tmab-8**, comprising PEG spacers before and after the TCO and chemical structure of trigger **10**.

The blood clearance of [^**89**^**Zr]Zr-Tmab-8** (0.5 mg/kg) alone in tumor-free mice was slow and bi-phasic (1.86 h t_1/2,α_ and 21.50 h t_1/2,β_), and matches the typical blood clearance of ^89^Zr-labeled Tmab, suggesting that the TCO-modification does not change the in vivo behavior of the radioimmunoconjugate (Figure 6A, Group A; Figure 6D).

**Figure 6.**
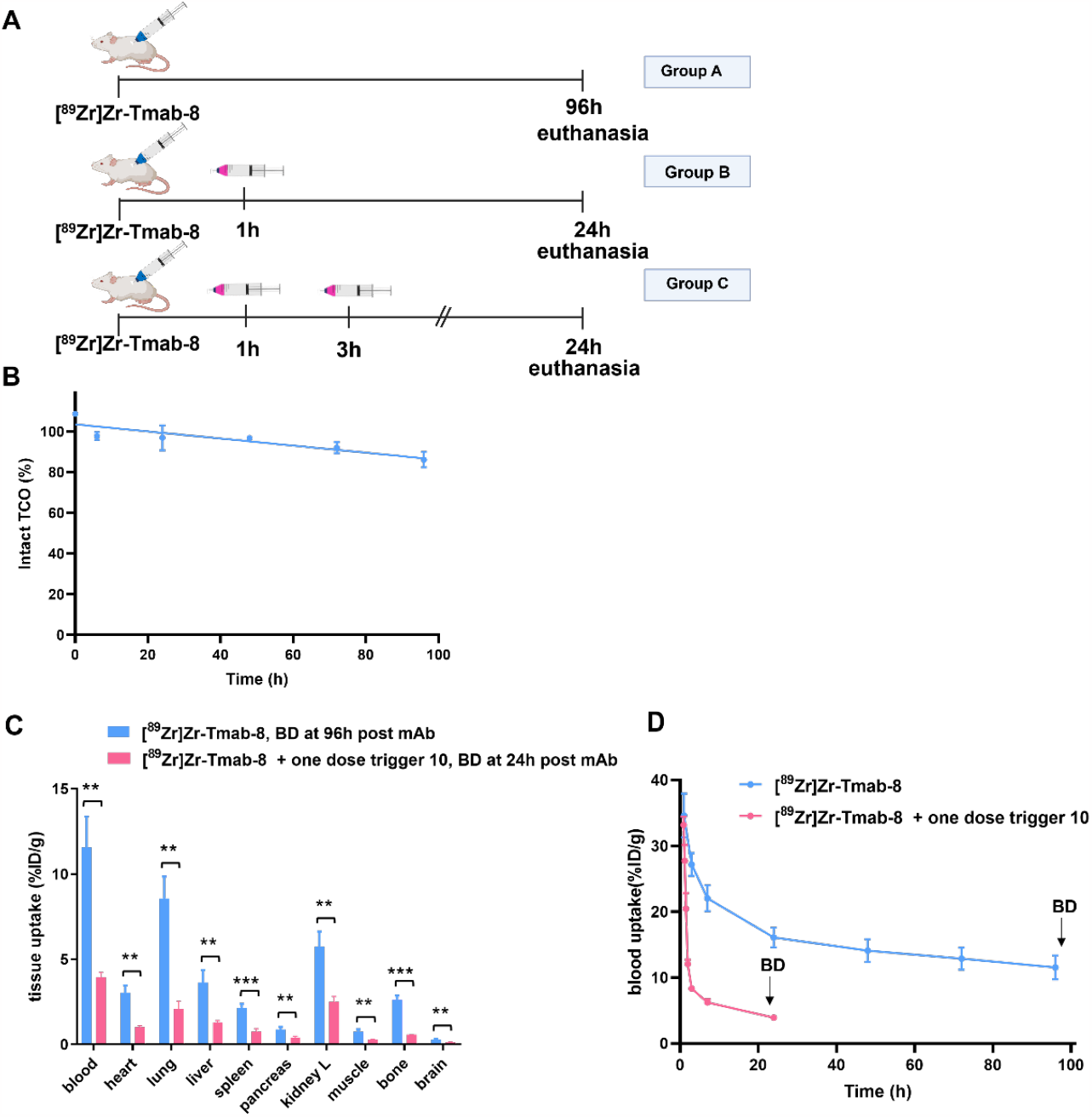
Evaluation of [^89^Zr]Zr-Tmab-8 and trigger 10 in tumor-free mice. **(A)** Experimental scheme of in vivo studies, where tumor-free mice received [^**89**^**Zr]Zr-Tmab-8** alone (Group A), mAb followed by one dose of trigger **10** (Group B) or two doses of trigger **10** (Group C). **(B)** Normalized in vivo TCO linker stability in tumor-free mice. The data represent the mean % ± SD (n=4). (**C)** Biodistribution at respectively 96 h and 24 h post-mAb injection of mice treated with [^**89**^**Zr]Zr-Tmab-8** alone (blue) and in combination with one dose of trigger **10** 1 h post-[^**89**^**Zr]Zr-Tmab-8** (pink). The data represent the mean % ± SD (n=4). Statistical analysis was performed using the unpaired Student’s t-test, **p < 0.01 and ***p < 0.001). **(D)** Rapid removal of radioactivity in mice receiving trigger **10** 1 h post-[^**89**^**Zr]Zr-Tmab-8** administration. The data represent the mean % ± SD (n=4).

Regarding the triggered release, previous studies have shown that the radioactive moiety [^89^Zr]Zr-DFO is eliminated from the blood via the kidneys within 20-60 min and we assumed that the released fragment (Figure 5) from circulating [^**89**^**Zr]Zr-Tmab-8** would exhibit similar pharmacokinetics [32,33]. We were therefore pleased to observe that in mice that received [^**89**^**Zr]Zr-Tmab-8** followed by one dose of trigger **10** (33.4 μmol/kg) (Figure 6A, Group B) the radioactivity in blood was 4-fold lower compared to the mice that did not receive the trigger, with 3.93 ± 0.29% ID/g and 16.08 ± 1.51% ID/g (p=0.0004) at 24 h post-mAb injection, respectively. This reduction corresponds well with the in vitro cleavage yield but to check if there was any unreacted TCO left in blood, a second dose of trigger **10** was administered 2 h after the first dose (Figure 6A, Group C). No significant decrease of the radioactivity in blood (3.61 ± 0.20 % ID/g, p=0.1291) was observed after the second trigger dose, confirming that all TCO in blood had already reacted with the first trigger dose (Table S1, Figure S6).

Previous studies have determined that TCO can isomerize to the unreactive *cis*-isomer in the presence of serum proteins [34]. To assess the deactivation rate of the TCO as Tmab-DFO linker we collected blood samples from healthy mice that received [^**89**^**Zr]Zr-Tmab-8** and reacted those with an excess of **10** ex vivo followed by SEC analysis to quantify the fragment release percentage as a measure of TCO content. The in vivo TCO deactivation half-life of the [^**89**^**Zr]Zr-Tmab-8** was calculated to be 16.5 days (Figure S5), which is 3-fold higher than what we have found for TCO-linked ADCs [20,21], allowing long intervals between mAb and trigger if needed.

### Release experiments in cell culture and in tumor-bearing mice

The presented approach centers on the use of a non-cell-permeable trigger to release the radioactive fragment from the mAb in blood at a time point where enough radiolabeled mAb has internalized into cancer cells, out of reach of the trigger. To get an indication of the required interval between mAb and trigger injection we first investigated the binding and internalization of [^**89**^**Zr]Zr-Tmab-8** in HER2-positive BT-474 cancer cells, which we found to be fast. The total mAb cell binding increased from 74.2% to 91.3% from 6 h to 24 h, with only 17.4% and 9.9% of the radioactivity surface-bound at these two time points (Figure S4). These findings align with published data showing that indeed some trastuzumab remains on the cell surface [35], although in our studies this fraction is much smaller.

Then we set out to investigate the release of [^**89**^**Zr]Zr-Tmab-8** by trigger **10** in mice bearing orthotopic BT-474 cancer xenografts, with 6 h and 24 h interval between mAb and trigger administration (biodistribution at 4 h post-trigger; Figure 7, Table S2 and S3). When the trigger was administered 6 h post-mAb, the tumor-blood (T/B) ratios reached 2.3 ± 0.6, which is 2.3-fold higher than the T/B of 1.0 ± 0.4 for the control group, p=0.0057 (Figure 7B). The 24 h interval afforded a T/B ratio of 6.6 ± 0.9, which is 2.6-fold higher than the T/B of 2.5 ± 0.7 for the control group, p<0.0001 (Figure 7B). The slightly higher T/B ratio improvement at 24 h time point is consistent with the slightly higher internalization found in vitro in cells at that time and is due to the increased retention of radioactivity in the tumor at 24 h in the group receiving the trigger (Figure S4). Furthermore an improved tumor-organ ratio was observed for blood-rich organs, such as the liver (from 3.8 ± 1.1 to 6.0 ± 1.9, p=0.0673 for 6 h and from 9.3 ± 2.2 to 13.6 ± 3.4, p=0.0494 for 24 h) and lungs (from 2.2±0.8 to 4.0±1.0, p=0.0148, for 6 h and from 4.8±1.8 to 10.8±2.9, p=0.0066, for 24 h), after the trigger injection at both time points (Table S3 and Figure 7B). The complete intratumor retention of the radioactivity already at 24 h p.i. combined with the pronounced reduction in blood is promising for future clinical applications, as the Tmab blood clearance will be much slower in humans while the tumor cell internalization rate is likely to be the same, potentially affording a larger reduction of the radiation dose in blood.

**Figure 7.**
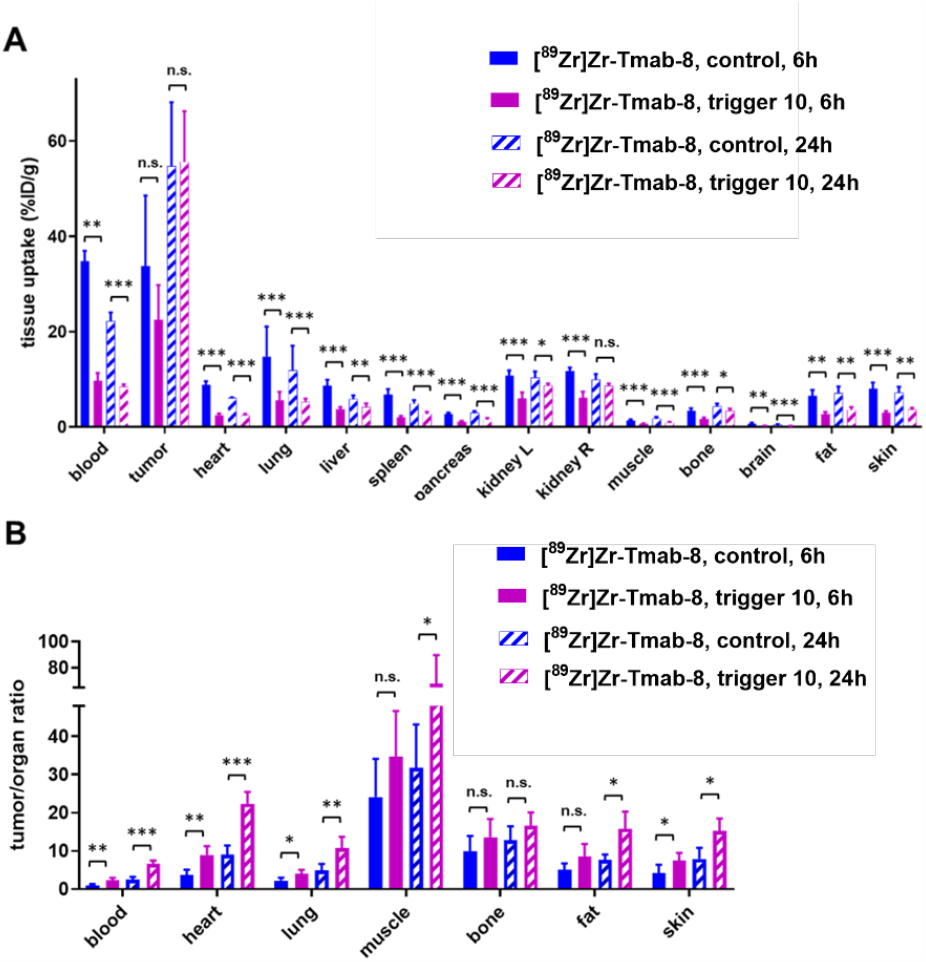
Evaluation of [^89^Zr]Zr-Tmab-8 and trigger 10 in mice bearing BT-474 xenografts at two different time points. **(A)** Biodistribution (4 h post-trigger administration) of mice bearing BT-474 xenografts receiving [^**89**^**Zr]Zr-Tmab-8** and trigger **10** at 6 h and 24 h post-mAb. The data represent the mean % ± SD (n=5). Statistical analysis was performed using the unpaired Student’s t-test, tumor 6 h (p=0.1793), tumor 24 h (p=0.9042). **(B)** Tumor-organ ratios for mice receiving [^**89**^**Zr]Zr-Tmab-8** and trigger **10** at 6 h and 24 h. The data represent the mean % ± SD (n=5). Statistical analysis was performed using the unpaired Student’s t-test, tumor-muscle 6 h (p=0.1682), tumor-bone 6 h (p=0.2119), tumor-fat 6 h p=0.0719), tumor-bone 24 h (p=0.1219). *p< 0.05, **p< 0.01, ***p < 0.001 and n.s.=not significant.

### Small animal PET imaging

Given the comparable T/B ratio improvement with the 6 h and 24 h interval we questioned whether the tumor-nontumor ratio improvement at 6 h would allow effective same-day imaging of a radiolabeled mAb. To this end, mice received [^**89**^**Zr]Zr-Tmab-8** followed by **10** 6 h later with PET scans taken 1 h prior and 4 h post-trigger administration (Figure 8, Table 2, Figure S7). The results clearly indicate that before the trigger administration (scan 1) there are high radioactivity levels in blood and in blood-rich organs (tumor-heart ratio 0.9 ± 0.1, p<0.0229), which increases the background and hinders the visibility of the tumor (red arrow, scan 1 in Figure 8B). To our delight, upon triggered chelate release the background radioactivity was largely reduced, greatly improving the tumor image (tumor-heart ratio 2.4 ± 0.7, p<0.0229, red arrow, scan 2 in Figure 8B). The PET images obtained are in accordance with the biodistribution data, which afforded a T/B ratio of 2.3 ± 0.6 (p<0.0057) and they show that the clearance of the released small fragment occurs via the kidneys into the bladder (yellow arrow, scan 2 in Figure 8B). In a pretargeting study using TCO-functionalized trastuzumab and a ^18^F-labeled tetrazine tracer, Lewis *c*.*s*. reported lower T/B ratios (0.57 ± 0.04), likely due to the low amount of non-internalized TCO-trastuzumab available for the tetrazine reaction at 48 h [36]. In contrast, the click-to-release approach presented herein uses the fast mAb internalization to its advantage.

**Figure 8.**
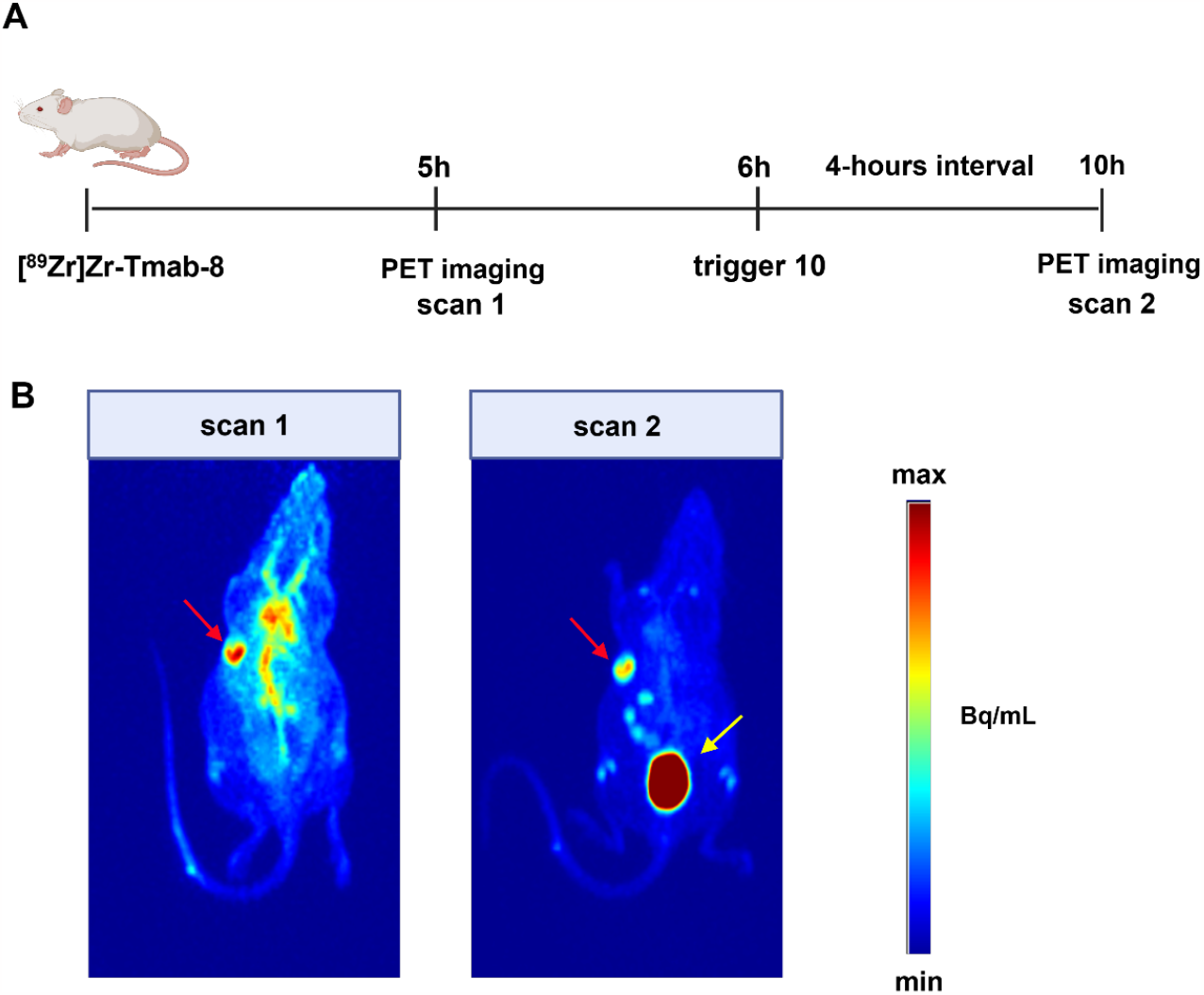
PET imaging studies in tumor-bearing mice. **(A)** Experimental scheme of PET imaging studies. Mice bearing HER2-positive BT-474 xenografts (n=4) received [^**89**^**Zr]Zr-Tmab-8** and 5 h later were imaged under anesthesia (scan 1). One hour later the mice received trigger **10** and 4 h post-trigger **10** administration were imaged again under anesthesia (scan 2) **(B)** Representative PET maximum intensity projections from the same mouse, before trigger administration (scan 1) and after trigger administration (scan 2). The images are scaled to the same min and max values. Red arrow indicates the tumor site and yellow arrow indicates the bladder.

**Table 2.** PET quantification. Mice (n = 4) bearing BT-474 xenografts were injected with [^**89**^**Zr]Zr-Tmab-8** (ca 0.5 mg/kg, ca 5 MBq) and 5 h later were imaged to obtain scan 1. After 1 h they received a dose of trigger **10** and 4 h post-trigger injection they were imaged again to obtain scan 2. PET scans were reconstructed and the tumor-muscle and tumor-heart ratios were obtained. Data were compared using unpaired t-test to obtain p=0.0229 for tumor-heart and p=0.1248 for tumor-muscle. The data represent the mean % ± SD (n=4).

| **PET scans** | **tumor-muscle ratio** |
| --- | --- |
| scan 1 | 8.4 ± 2.3 |
| scan 2 | 20.2 ± 8.8 |
|  | **tumor-heart ratio** |
| scan 1 | 0.8 ± 0.1 |
| scan 2 | 2.4 ± 0.7 |

## Conclusion

We have demonstrated the successful use of click-to-release chemistry for the efficient and temporally controlled cleavage of a radioactive chelator from an antibody in vivo. This approach provided excellent off-target deactivation of the radiolabeled mAb with fast renal clearance of the cleaved radioactive fragment while largely retaining the on-target activity in the tumor both when using the trigger at 6 h and 24 h post-mAb administration. Future work will focus on further increasing the in vivo cleavage yields to maximize the benefit. We believe that the presented click-cleavable radioimmunoimaging approach may allow for lower whole body radiation doses in the clinic, potentially same day imaging, and improved image quality. While the present report demonstrates the benefits in the context of a cancer cell-internalized mAb we expect the approach to be equally promising for imaging of mAbs designed to cross the blood brain barrier, allowing selective radioactive background reduction in cerebral blood. We think the approach may also find application in increasing target-nontarget ratios of smaller targeting agents. Finally, we believe it may be highly beneficial for radioimmunotherapy by reducing the bone marrow dose, thereby enabling increased tumor doses.

## Acknowledgements

We thank Martijn den Brok (Tagworks Pharmaceuticals), Gerben Fransen, Bianca Lemmers-van de Weem, Kitty Lemmens-Hermans and the staff at PRIME (Radboud University Medical Center) for assistance with the experiments. This work was financed by the European Union’s Horizon 2020 research and innovation program under the Marie Sklodowska-Curie Grant Agreement no. 765497 (THERACAT).

