## supplementary information for "Click-to-Release: Cleavable Radioimmunoimaging with ^89^Zr-DFO-*Trans*-Cyclooctene-Trastuzumab Increases Tumor-to-Blood Ratio"

### *Reagents*

All reagents and solvents were obtained from commercial sources (Sigma-Aldrich, Acros, Merck) and used without further purification, unless stated otherwise. 1-Amino-3,6,9,12-tetraoxapentadecan-15-oic-acid and N-(29-amino-3,6,9,12,15,18,21,24,27-nonaoxanonacosyl)-3-(2,5-dioxo-2,5-dihydro-1H-pyrrol-1-yl)propenamide as a TFA salt were purchased from Broadpharm. Tris(2-Carboxylethyl)phosphine (TCEP) was purchased from Thermo Fisher Scientific.

Trastuzumab solutions were purchased from Mylan (Ogivri) and were reconstituted following the manufacturer's instructions. Trastuzumab was purified using PD-10 cartridges (Cytiva) eluted with PBS. The concentration of the collected vials was determined by Nanodrop and the solutions were stored at -80 ºC. [$^{111}$In]Indium chloride and [$^{89}$Zr]zirconium oxalate were purchased from Curium Pharma and Perkin Elmer, respectively. Water was distilled and deionized (18 MΩcm) by means of a milliQ-water filtration system (Millipore). Sterile phosphate buffered saline (PBS) was purchased from Fresenium Kabi. Amicon Ultra centrifugal devices (30kDa MW cut-off) were purchased from Millipore. Mouse serum was purchased from Innovative Research and was filtered through 0.2 μm filters before use. Zeba desalting spin columns (40kDa MW cut-off, 0.5mL) were purchased from Thermo Fisher Scientific. Chelex 100 (200-400 mesh) was purchased from Bio-Rad. For animal experiments, matrigel was purchased from Corning Life Sciences, the BT-474 cancer cell line was purchased from ATCC and 17β-estradiol releasing pellets (0.18 mg, 60 days release) were purchased from Innovative Research of America.

### *Instrumentation*

NMR characterization of compounds was carried out on a Bruker AVANCE HD nanobay console with a 9.4 T Ascend magnet (400 MHz) and a Bruker AVANCE III console with a 11.7 T UltraShield Plus magnet (500 MHz) equipped with a Bruker Prodigy cryoprobe. Chemical shifts are reported in ppm downfield from TMS at 25 ºC. Abbreviations used from splitting patterns are s=singlet, t=triplet, q-quartet, m=multiplet and br=broad. Reverse phase (RP) liquid chromatography was performed on a Shimadzu HPLC system with MeCN/water mixtures (containing 0.1% TFA) as the eluent. LC-MS was recorded using Thermo Finigan LCQ Fleet system, applying a gradient of water and MeCN containing 0.1% TFA. Size exclusion Chromatography (SEC) was carried out on an AKTA-purifier system (Cytiva) equipped with a UV detector, a Gabi radioactive detector and a fraction collector. The samples were loaded on a Superdex200 10/300 column (Cytiva) which was eluted with PBS with a flow rate of 0.6 mL/min. Radio-TLC was performed on ITLC-SG strips obtained by Agilent Technologies and eluted with 0.1M sodium citrate pH 6 and imaged on a phosphor imager (Typhoon FLA 7000; Cytiva). In these conditions, free $^{89}Zr$ migrates with $R_f$=0.9, while $^{89}Zr$-labeled mAb remains at the origin. UV measurements were carried out on a Tecan Infinite 200 microplate reader. The antibody concentrations were measured using a Nanodrop 1000 spectrometer (Thermo Fisher Scientific) at 280 nm using a program for IgGs.

### *Synthesis of linker-chelator 2*

**2,5-Dioxopyrrolidin-1-yl(1R,6R,E)-1-methyl-6-(((3,14,25-trihydroxy-2,10,13,21,24-pentaoxo-3,9,14,20,25-pentaazatriacontan-30-yl)carbamoyl)oxy)cyclooct-4-ene-1-carboxylate (2).**

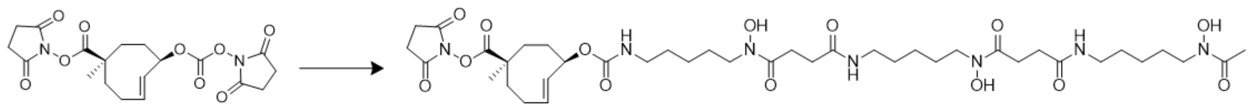

1 2

2,5-Dioxopyrrolidin-1-yl-6-((((2,5-dioxopyrrolidin-1-yl)oxy)carbonyl)oxy)-1-methylcyclooct-4-ene-1-carboxylate **1** was synthesized according to the literature procedure [1]. A mixture of **1** (15 mg, 0.035 mmol) and deferoxamine mesylate salt (29.9 mg, 0.046 mmol) in DMSO (1 mL) were stirred for 4 h at RT. The reaction was monitored by LC-MS. Upon completion of the reaction, the reaction mixture was diluted with water containing 0.1% TFA, followed by preparative RP-HPLC purification, using an elution gradient of 5% to 95% MeCN in water (both containing 0.1% TFA) to yield compound **2** (25.7 mg, 0.029 mmol, 84%) after lyophilization as a fluffy white powder. $^1H$ NMR (400 MHz, $CDCl_3$) δ 5.87 (m, 1H), 5.63 (m, 1H), 5.17 (s, 1H), 3.64 (s, 3H), 3.62 (s, 1H), 3.21 (s, 4H), 2.82 (s, 4H), 2.67 (m, 4H), 2.29 (m, 3H), 2.23 (m, 3H), 2.08 (m, 2H), 1.95 (m, 2H), 1.84 (m, 8H), 1.65 (m, 4H), 1.54 (m, 5H), 1.42 (m, 5H), 1.26 (s, 3H),1.19 (m, 3H) ppm. HPLC-MS/PDA: *m/z* =868.44 $[M+H]_+$, calcd. 867.46 for $C_{40}H_{65}N_7O_{14}$.

### *Synthesis of linker-chelator 4*

**2,5-Dioxopyrrolidin-1-yl(1R,6R,E)-1-methyl-6-((methyl(9,20,31-trihydroxy-2,10,13,21,24,32-hexaoxo-3,9,14,20,25,31-hexaazatritriacontyl)carbamoyl)oxy)cyclooct-4-ene-1-carboxylate (4).**

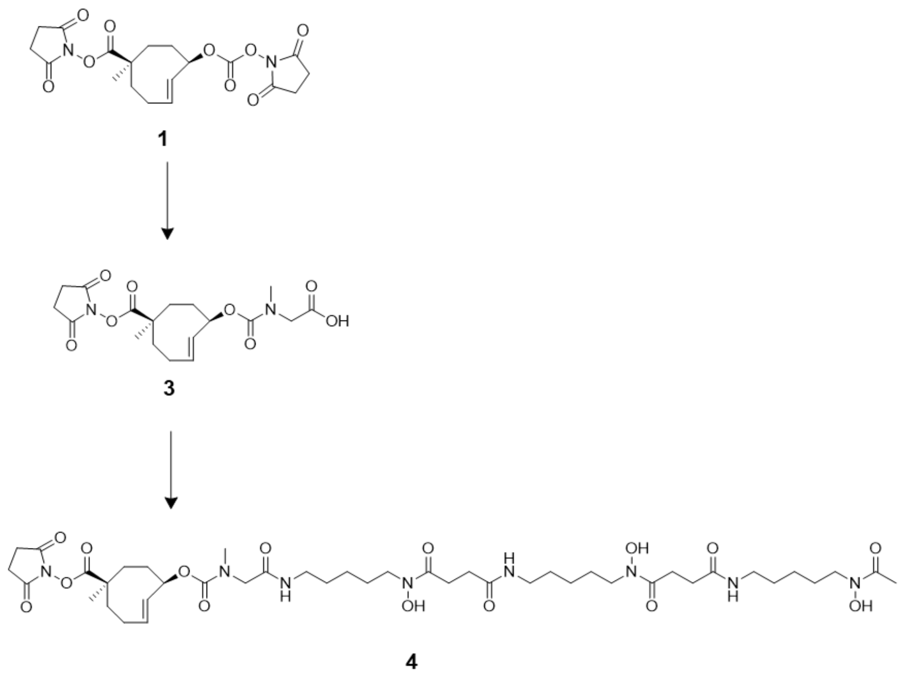

Compound **1** (15 mg, 0.035 mmol) and sarcosine (3.1 mg, 0.035 mmol) were mixed in water for 2 h. Water was removed and the formed product **3** was combined, without further purification, with PyBOP (18.2 mg, 0.035 mmol) and DIPEA (12.1 µL, 0.07 mmol) in DMSO and the reaction mixture was stirred in RT for 10 min, before adding deferoxamine mesylate salt (24.9 mg, 0.038 mmol). The reaction mixture was stirred in RT for 3 h and was monitored by LC-MS. Upon completion of the reaction, the reaction mixture was diluted with water containing 0.1% TFA, followed by preparative RP-HPLC purification, using an elution gradient of 5% to 95% MeCN in water (both containing 0.1% TFA) to yield compound **4** (28 mg, 0.029 mmol, 83%) after lyophilization as a fluffy white powder.$^{1}$H NMR $^{1}$H NMR (400 MHz, DMSO-$d_6$) δ 5,77 (m, 1H), 5.09 (m, 1H), 3.84 (m, 1H), 3.40 (m, 22H), 3.04-2.80 (m, 6H), 2.59-2.54 (m, 2H), 2.27-2.22 (m, 3H), 1.95 (s, 2H), .89-1.76 (m, 2H), 1.50-1.32 (m, 5H), 1.22 (m, 4H) ppm. HPLC-MS/PDA: *m/z* =939.20 $[M+H]_+$, calcd. 938.50 for $C_{43}H_{70}N_8O_{15}$.

### *Synthesis of linker-chelator 6*

**2,5-Dioxopyrrolidin-1-yl(1R,6R,E)-1-methyl-6-((methyl(25,36,47-trihydroxy-2,18,26,29,37,40,48-heptaoxo-6,9,12,15-tetraoxa-3,19,25,30,36,41,47-heptaazanonatetracontyl)carbamoyl)oxy)cyclooct-4-ene-1-carboxylate (6).**

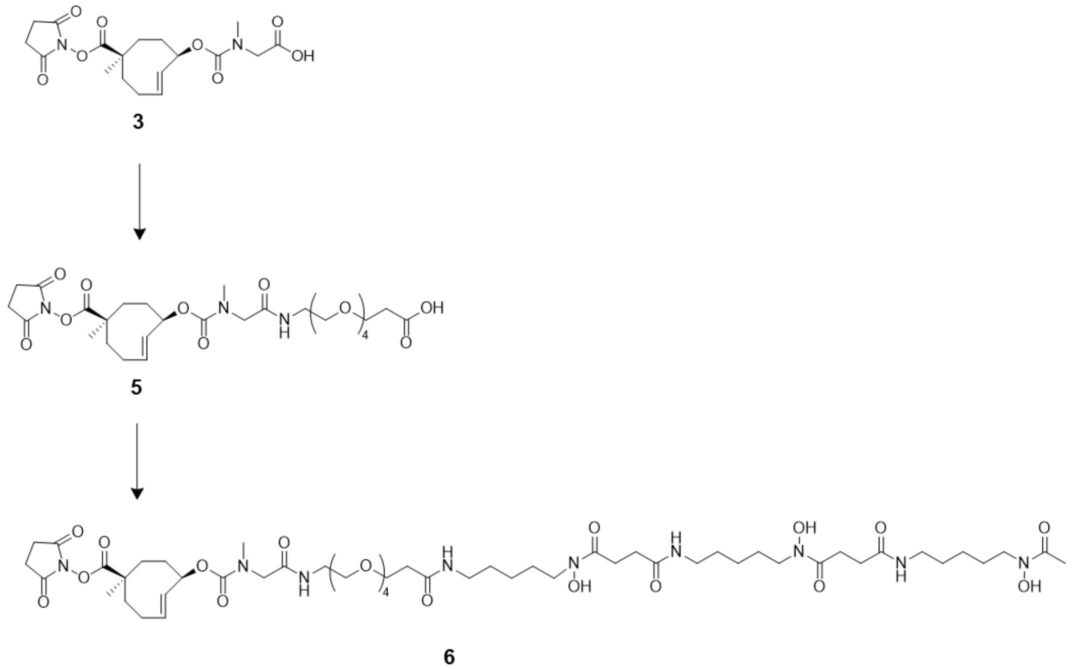

PyBOP (19.8 mg, 0.038 mmol) and DIPEA (13.2 µL, 0.076 mmol) were added to a stirred solution of **3** (15 mg, 0.038 mmol) in DMF at RT for 10 min. Upon activation, a solution of amino-PEG4-acid (12.2 mg, 0.046 mmol) and DIPEA (17.79 uL, 0.102 mmol) was added, and the solution was stirred for 4 h. Upon completion of the reaction, the reaction mixture was diluted with acidified water containing 0.1% TFA, followed by preparative RP-HPLC purification, using an elution gradient of 5% to 95% MeCN in water (both containing 0.1% TFA) to yield compound **5** (20 mg, 0.031 mmol, 82%) after lyophilization as a fluffy white powder.

PyBOP (16.1 mg, 0.031 mmol) and DIPEA (10.8 µL, 0.062 mmol) were added to a solution of **5** (20 mg, 0.031 mmol) in DMSO and the reaction was stirred at RT for 10 min. A solution of deferoxamine mesylate salt (22.3 mg, 0.034 mmol) and DIPEA (10.8 µL, 0.062 mmol) in DMSO was added and the reaction mixture was monitored by LC-MS. After 4 h, the reaction mixture was diluted with water containing 0.1% TFA, followed by preparative RP-HPLC purification, using an elution gradient of 5% to 95% MeCN in water (both containing 0.1%

TFA) to yield compound **6** (32 mg, 0.026 mmol, 86 %) after lyophilization as a fluffy white powder. $^{1}$H NMR (400 MHz, $CDCl_3$) δ 5.88 (m, 1H), 5.66 (m, .1H), 5.24 (s, 1H), 3.89 (m, 6H), 3.77 (m, 2H), 3.64 (m, 15H), 3.49 (m, 2H), 3.06 (m, 3H), 2.82 (s, 3H), 2.60 (m, 2H), 2.38 (m, 1H), 2.30 (m, 2H), 2.15 (m, 3H), 1.92 (m, 2H), 1.27 (m, 3H) ppm. HPLC-MS/PDA: *m/z* =1186.32 $[M+H]^+$, calcd. 1185.64 for $C_{54}H_{91}N_9O_{20}$.

### *Synthesis of linker-chelator 7*

**(1R,6R,E)-6-((33-(2,5-Dioxo-2,5-dihydro-1H-pyrrol-1-yl)-31-oxo-3,6,9,12,15,18,21,24,27-nonaoxa-30-azatritriacontyl)carbamoyl)-6-methylcyclooct-2-en-1-yl(3,14,25-trihydroxy-2,10,13,21,24-pentaoxo-3,9,14,20,25-pentaazatriacontan-30-yl)carbamate (7).**

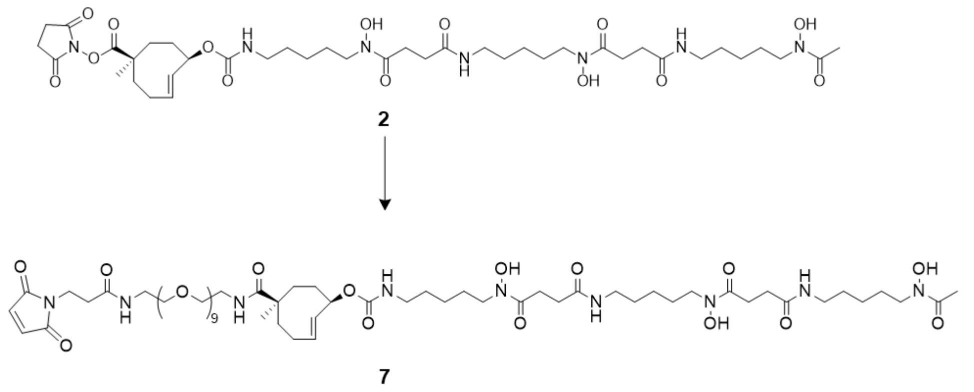

A mixture of maleimido-PEG9-amine TFA salt (14 mg, 0.023 mmol) in DMF (0.5 mL) and DIPEA (16.2 µL, 0.092 mmol) was added to a solution of **2** (20 mg, 0.023 mmol) in DMF (0.5 mL). The reaction mixture was stirred at RT and monitored by LC-MS. Upon the formation of the desired product, the reaction mixture was diluted with water containing 0.1% TFA, followed by preparative RP-HPLC purification, using an isocratic elution of 20% MeCN in water (both containing 0.1% TFA) to yield compound **7** (4.5 mg, 0.003 mmol, 14%) after lyophilization as a colorless oil. $^{1}$H NMR (500 MHz, $CDCl_3$) δ 6.70 (s, 2H), 5.87 (m, 1H), 5.19 (s, 1H), 3.83 (m, 1H), 3.65-3.61 (m, 17H), 3.42 (m, 2H), 3.21 (m, 2H), 2.80-2.62 (m, 3H), 2.53 (m, 1H), 2.29-2.23 (m, 2H), 2.11-1.91 (m, 2H), 1.17 (m, 13H). 1.37 (m, 4H), 1.12 (m,

2H), 0.95-0.83 (m, 1H) ppm. HPLC-MS/PDA: *m/z* =1360.40 $[M+H]^+$, calcd. 1359.76 for $C_{63}H_{109}N_9O_{23}$.

### *Synthesis of linker-chelator 8*

**(1R,6R,E)-6-((33-(2,5-Dioxo-2,5-dihydro-1H-pyrrol-1-yl)-31-oxo-3,6,9,12,15,18,21,24,27-nonaoxa-30-azatritriacontyl)carbamoyl)-6-methylcyclooct-2-en-1-yl methyl(25,36,47-trihydroxy-2,18,26,29,37,40,48-heptaoxo-6,9,12,15-tetraoxa-3,19,25,30,36,41,47-heptaazanonatetracontyl)carbamate (8).**

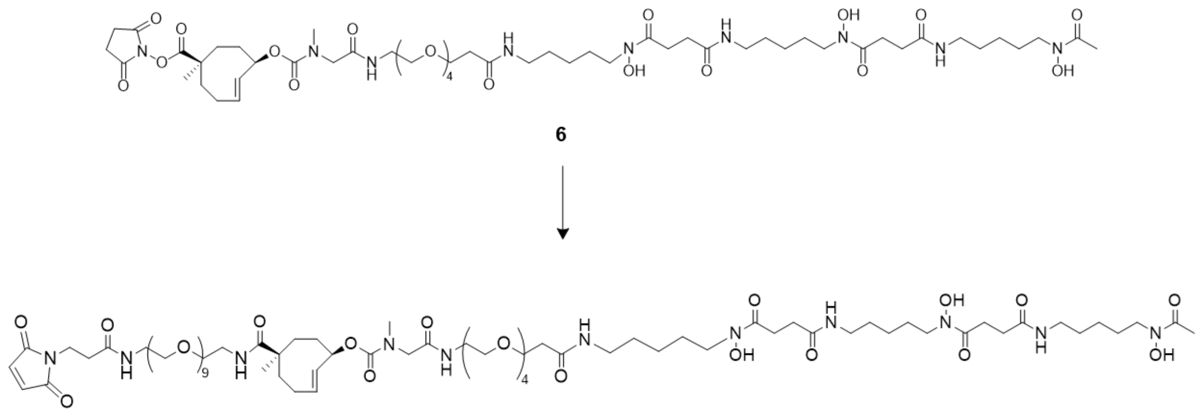

8

A mixture of maleimido-PEG9-amine TFA salt (4 mg, 0.023 mmol) in DMF (0.5 mL) and DIPEA (6.2 µL, 0.092 mmol) was added to a solution of **6** (20 mg, 0.023 mmol) in DMF (0.5 mL). The reaction mixture was stirred at RT and monitored by LC-MS. Upon the formation of the desired product, the reaction mixture was diluted with water containing 0.1% TFA, followed by preparative RP-HPLC purification, using an isocratic elution of 20% MeCN in water (both containing 0.1% TFA) to yield compound **8** (4.2 mg, 0.002 mmol, 11 %) after lyophilization as a colorless oil. Isolated compound **8** contains an impurity resulting from addition of maleimido-PEG9-amine to the maleimide moiety in **8** (ca. 10% by analytical HPLC). $^1$H NMR (500 MHz, $CDCl_3$) δ 6.70 (s, 2H), 5.87 (m, 1H), 5.62 (m, 1H), 5.19 (s, 1H), 3.99-3.94 (m, 1H). 3.84 (m, 1H), 3.72 (m, 1H), 3.65-3.61 (m, 17 H), 3.53 (m, 2H), 3.49-3.46 (m, 1H), 3.41 (m, 1H), 3.21 (m, 3H), 3.08-3.01 (m, 2H), 2.83 (m, 2H), 2.62-2.57 (m, 2H), 2.53-2.49 (m, 2H), 2.04-2.01 (m, 1H), 1.31-1.19 (m, 21H), 1.12 (m, 4H), 0.89-0.81 (m, 7H) ppm. HPLC-MS/PDA: *m/z* =1677.56 $[M-H]^+$, calcd. 1677.94 for $C_{77}H_{135}N_{11}O_{29}$.

### *Preparation of conjugates Tmab-2, Tmab-4, Tmab-6, Tmab-7, Tmab-8*

Trastuzumab was conjugated to compounds **2**, **4**, **6** via NHS chemistry. Typically, trastuzumab (1 mg) was reacted for 2 h with the linker-chelator (35 eq) in PBS (final volume 4 mg/mL, pH adjusted to ca 8.8 with 1M sodium carbonate). The crude reaction mixture was purified using SEC chromatography and chelex-treated PBS as an eluent and then the purified mAb conjugate was stored in aliquots at -70 ºC for further use. Typically, this procedure afforded ca 1.6 linker-chelator per antibody, as determined by a tetrazine titration with $^{111}$In-labeled tetrazine, analyzed by SDS-PAGE [1].

Trastuzumab was conjugated to compounds **7** and **8** via maleimide chemistry using a modification of an already published procedure [2]. Trastuzumab (1 mg) was partially reduced with 2.3 eq TCEP in PBS at 37 ºC for 30 min. Then, the solution was 1:1 diluted with PBS containing 5mM EDTA (pH adjusted to 6.8) and was cooled on ice. The reduced antibody was reacted with the linker-chelator **7** or **8** (10 eq, 10 mg/mL in DMSO) for 30 min on ice and then overnight at 4 ºC in the dark. The crude reaction mixture was purified by SEC using chelex-treated PBS as eluent and then stored in aliquots in -70 ˚C for further use. Typically, this procedure afforded 2.5 linker-chelators per antibody, as measured by tetrazine titration [1].

### *Binding and internalization assay*

The HER2 positive BT-474 cells were cultured in RPMI medium supplemented with 2 mM glutamine and 10% fetal calf serum. Approximately 48 h prior to the experiment, the cells were plated in 6-well plates at 0.6 million cells/well in 3 mL medium. At the time of the experiment, the cells were washed one time with pre-warmed PBS, followed by incubation for 6 or 24 h with 0.26 µg of $^{89}$Zr-conjugate in 3 mL binding medium (RPMI containing 0.5% BSA). Three wells were used for each condition. Blocking experiments were performed by adding a large excess (1000 eq) of trastuzumab to the medium. After incubation, the medium was removed and the cells were washed twice with ice-cold PBS followed by lysis in 0.1 M NaOH. To calculate the membrane bound activity, the cells were incubated with an acid buffer (0.1 M acetic acid, 154 mM NaCl, pH 2.6) on ice for 10 min. The lysates and acid wash solutions were measured by γ-counting together with standards to calculate the 100% added activity.

#### *Statistical analysis*

Group variation is described as the mean ± one standard deviation. GraphPad Prism version 9 was used for all statistical calculations.

### *Supplementary figures*

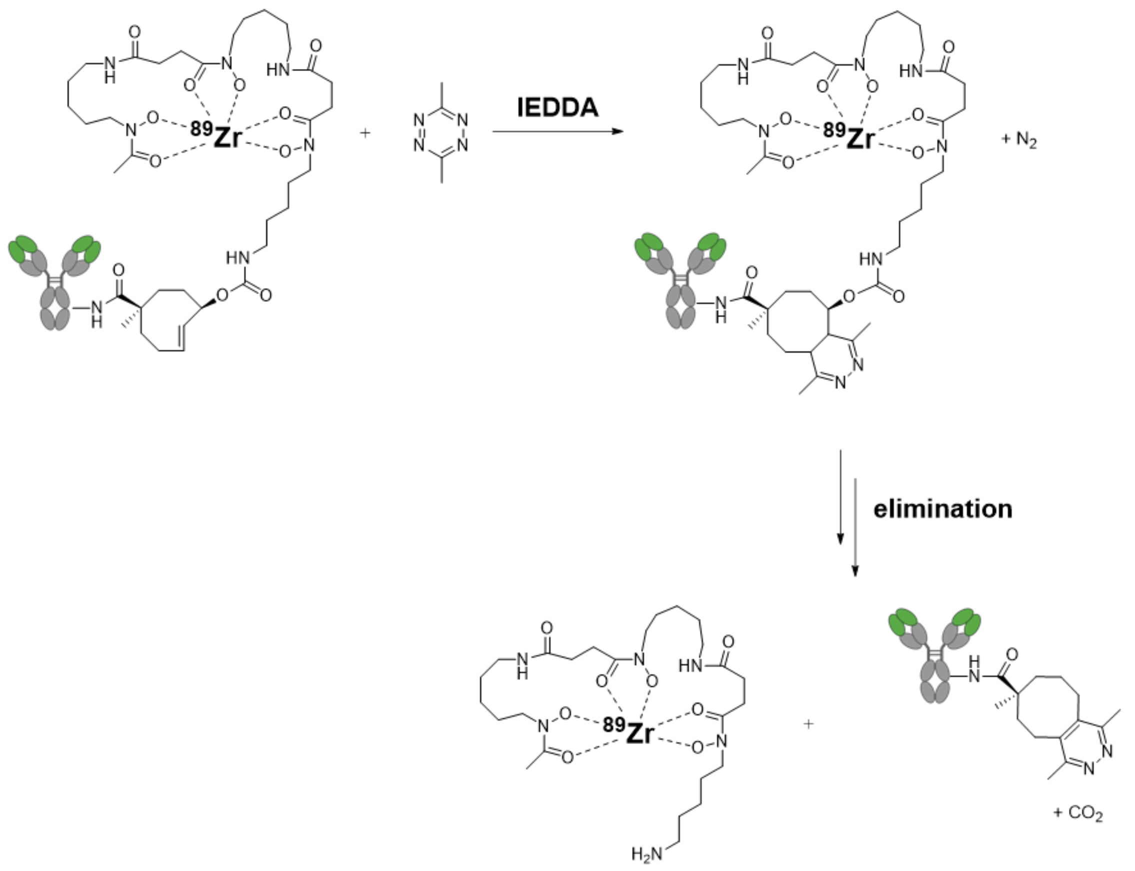

**Figure S1. Example of click-to-release reaction between conjugate [$^{89}$Zr]Zr-Tmab-2 and trigger 9.**

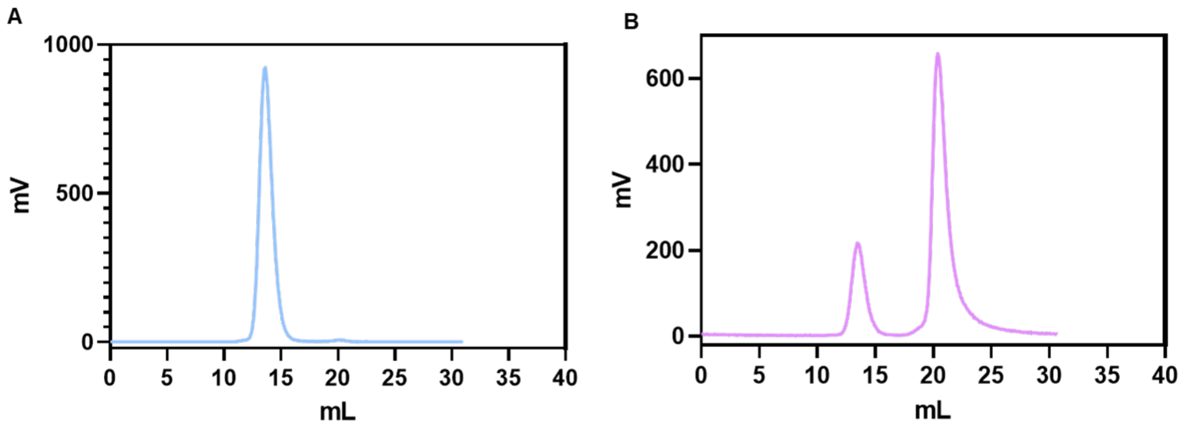

**Figure S2. Release experiments in PBS.** (A) SEC profile of **[$^{89}$Zr]Zr-Tmab-8** incubation overnight as a control and (B) SEC profile of **[$^{89}$Zr]Zr-Tmab-8** when reacted with 300 eq of trigger **10** in PBS for 24 h at 37 ºC.

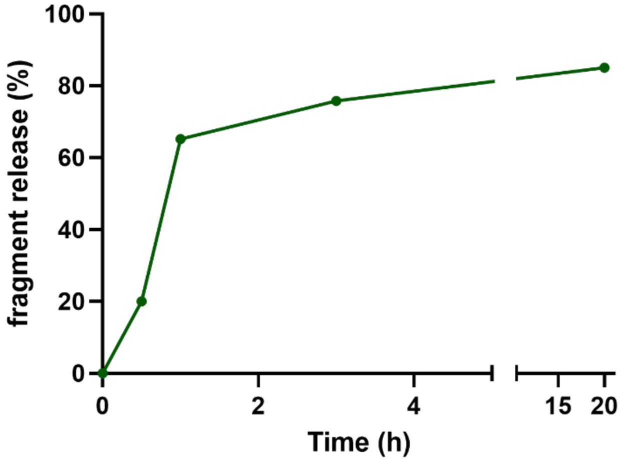

**Figure S3. Release experiments in plasma between [$^{89}$Zr]Zr-Tmab-8 and trigger 10.** The conjugate was radiolabeled with Zr-89 and was incubated with **10** (300 eq) in 50% mouse plasma in PBS at 37 ºC for up to 20 h. The kinetics of the release was monitored by SEC at different time points. Data are expressed as fragment release % (n=1).

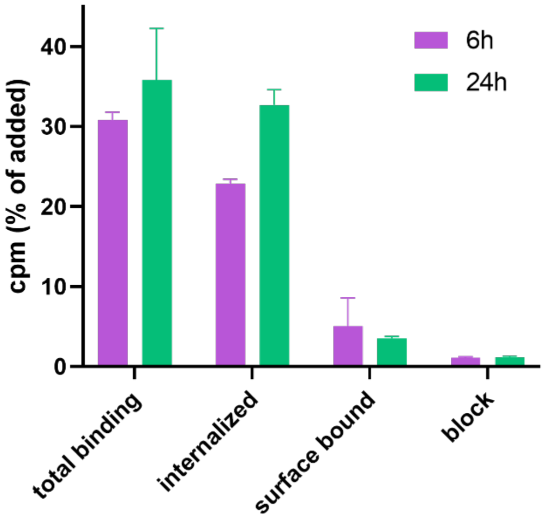

**Figure S4. Binding and internalization cell assay.** Total cell associated activity of cells expressed in % of added radioactivity. Cells were incubated with **[$^{89}$Zr]Zr-Tmab-8** for 6 h and 24 h. The bound and internalized radioactivity (total binding), the internalized radioactivity and the cell surface bound radioactivity were measured. Blocking was performed with 1000 eq of non-radiolabeled trastuzumab. Data are the mean with SD (n = 3).

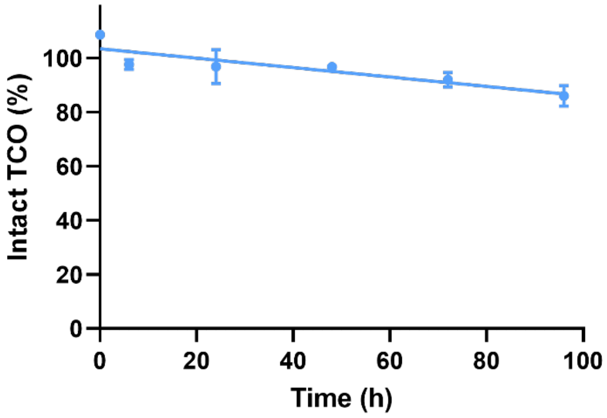

**Figure S5. Normalized in vivo linker stability of [$^{89}$Zr]Zr-Tmab-8.** Tumor-free nude mice (n=4) were injected with **[$^{89}$Zr]Zr-Tmab-8**. At selected time points (1 h, 3 h, 6 h 24 h, 48 h, 72 h and 96 h) blood samples were withdrawn. The blood samples reacted ex vivo with an excess of trigger **10** at 37 °C overnight in the dark followed by SEC analysis. The data points are the mean % release observed ± SD (n = 4) normalized to t=0.

.

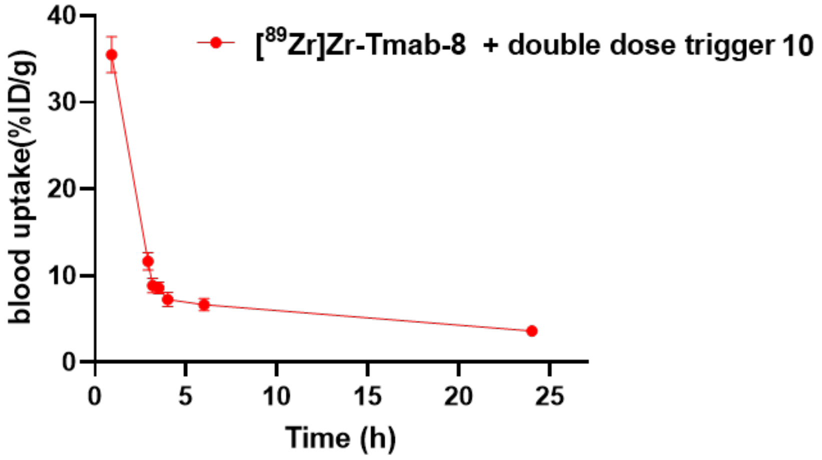

**Figure S6. Triggered release in tumor-free nude mice.** Blood kinetic studies in mice that received **[$^{89}$Zr]Zr-Tmab-8**. One hour post mAb injection the mice received a dose of trigger **10** (33.4 µmol/kg) and 2 h later the mice received an extra dose of trigger **10** (33.4 µmol/kg). The data points are the mean %ID/g ± SD (n = 4).

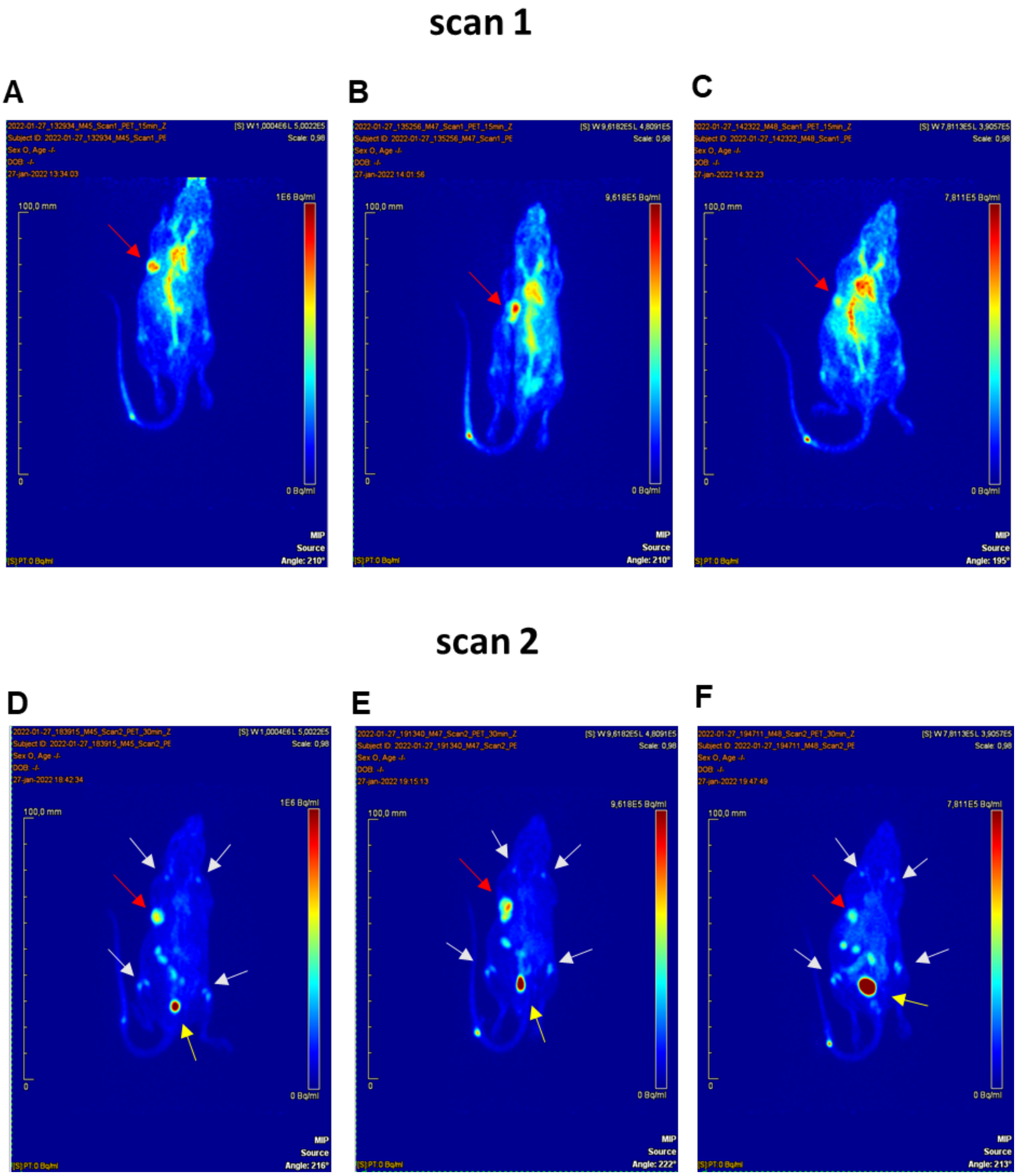

**Figure S7. PET imaging studies.** Mice were injected with **[$^{89}$Zr]Zr-Tmab-8** (ca. 0.5 mg/kg; 5 MBq in 100 µL) and 5 h later they were imaged under anesthesia obtaining scan 1 (A, B, C). One hour later, after recovery from anesthesia, the same mice received one dose of trigger **10** (33.4 µmol/kg) and 4 h post-trigger injection they were imaged again under anesthesia obtaining scan 2 (D, E, F). Images are presented as maximum intensity projections (MIPs), maximum intensity 9.62 $10^{-5}$ Bq/mL. In all images tumor site is indicated by a red arrow, bladder is indicated by a yellow arrow and the joints are indicated by white arrows.

### *Supplementary tables*

**Table S1.** Biodistribution studies in tumor-free mice. One group of mice received **[$^{89}$Zr]Zr-Tmab-8** (ca. 0.5 mg/kg, ca 0.5 MB in 100 µL) and 4 days later were euthanized (no trigger). One group of mice received **[$^{89}$Zr]Zr-Tmab-8** and 1 h post-mAb injection received trigger **10** (single dose trigger **10**; 33.4 µmol/kg). One group of mice received **[$^{89}$Zr]Zr-Tmab-8** and 1 h post-mAb injection received trigger **10** and 1 h later they received an extra dose of trigger **10** (double dose trigger **10**). Mice receiving trigger doses were euthanized 24 h after the last trigger injection. Data are the mean % ID/g with SD (n = 4).

| Organ (%ID/g) | no trigger | single dose trigger 10 | double dose trigger 10 |
|---|---|---|---|
| blood | 11.55 ± 1.82 | 3.93 ± 0.29 | 3.61 ± 0.20 |
| heart | 3.03 ± 0.45 | 1.01 ± 0.09 | 0.91 ± 0.07 |
| lung | 8.56 ± 1.29 | 2.09 ± 0.45 | 1.93 ± 0.46 |
| liver | 3.65 ± 0.72 | 1.28 ± 0.12 | 1.32 ± 0.17 |
| spleen | 2.16 ± 0.23 | 0.76 ± 0.16 | 0.85 ± 0.07 |
| pancreas | 0.88 ± 0.16 | 0.41 ± 0.07 | 0.29 ± 0.04 |
| kidney left | 5.73 ± 0.89 | 2.51 ± 0.32 | 2.67 ± 0.13 |
| kidney right | 5.87 ± 0.80 | 2.51 ± 0.14 | 2.89 ± 0.12 |
| muscle | 0.76 ± 0.13 | 0.28 ± 0.01 | 0.33 ± 0.17 |
| bone | 2.63 ± 0.24 | 0.57 ± 0.01 | 0.60 ± 0.06 |
| brain | 0.28 ± 0.04 | 0.12 ± 0.02 | 0.09 ± 0.01 |
| stomach* | 0.23 ± 0.04 | 0.12 ±0.05 | 0.09 ± 0.01 |
| small intestine* | 1.43 ± 0.11 | 0.50 ± 0.02 | 3.61 ± 0.20 |
| large intestine* | 0.61 ± 0.15 | 0.26 ± 0.07 | 0.91 ± 0.07 |

*These values are expressed in %ID/organ.

**Table S2.** Biodistribution studies in tumor-bearing mice. Two groups of mice received **[$^{89}$Zr]Zr-Tmab-8** (ca. 0.5 mg/kg, ca. 0.5 MB in 100 µL) and trigger **10** (33.4 µmol/kg) 6 h or 24 h post-mAb administration. Mice were euthanized 4 h after the trigger dose. Control mice that did not receive a trigger were euthanized 6 h and 24 h post-mb injection. Data are the mean % ID/g with SD (n =5).

| **Organ (%ID/g)** | **6 h no trigger** | **6 h trigger 10** | **24 h no trigger** | **24 h trigger 10** |
|---|---|---|---|---|
| blood | 34.78 ± 2.16 | 9.73 ± 1.56 | 22.21 ± 1.84 | 8.40 ± 0.50 |
| tumor | 33.75 ± 14.82 | 22.49 ± 7.24 | 54.68 ± 13.43 | 55.63 ± 10.60 |
| heart | 8.80 ± 0.79 | 2.48 ± 0.32 | 6.06 ± 0.14 | 2.47 ± 0.25 |
| lung | 14.79 ± 6.26 | 5.61 ± 1.73 | 11.91 ± 5.16 | 5.28 ± 0.68 |
| liver | 8.69 ± 1.22 | 3.77 ± 0.43 | 5.90 ± 0.75 | 4.19 ± 0.76 |
| spleen | 6.75 ± 1.23 | 2.10 ± 0.27 | 4.96 ± 0.70 | 2.76 ± 0.45 |
| pancreas | 2.75 ± 0.26 | 1,07 ± 0.17 | 2.96 ± 0.35 | 1.54 ± 0.35 |
| kidney left | 10.84 ± 1.05 | 6.02 ± 1.25 | 10.39 ± 1.24 | 8.71 ± 0.40 |
| kidney right | 11.71 ± 0.75 | 6.08 ± 1.33 | 9.91 ± 1.19 | 8.63 ± 0.47 |
| muscle | 1.40 ± 0.16 | 0.65 ± 0.12 | 1.79 ± 0.30 | 0.86 ± 0.18 |
| bone | 3.44 ± 0.55 | 1.68 ± 0.33 | 4.31 ± 0.62 | 3.38 ± 0.46 |
| brain | 0.75 ± 0.16 | 0.26 ± 0.06 | 0.52 ± 0.05 | 0.25 ± 0.02 |
| fat | 6.53 ± 1.24 | 2.67 ± 0.45 | 7.13 ± 1.38 | 3.62 ± 0.55 |
| skin | 8.03 ± 1.33 | 2.99 ± 0.39 | 7.21 ± 1.24 | 3.67 ± 0.39 |
| stomach* | 0.86 ± 0.18 | 2.01 ± 3.53 | 0.83 ± 0.15 | 0.98 ± 1.20 |
| small intestine* | 5.00 ± 0.41 | 4.88 ± 0.69 | 3.51 ± 0.45 | 4.31 ± 1.12 |
| large intestine* | 2.52 ± 0.19 | 7.58 ± 1.90 | 1.77 ± 0.59 | 7.06 ± 5.82 |

*These values are expressed in %ID/organ.

**Table S3.** Biodistribution studies (tumor-to-organ) in tumor-bearing mice. Mice received **[$^{89}$Zr]Zr-Tmab-8** (ca 0.5 mg/kg, ca 0.5 MBq in 100 µL) and trigger **10** (33.4 µmol/kg) 6 h and 24 h post-mAb administration. Mice were euthanized 4 h after the trigger dose in both cases. Control mice that did not receive a trigger were euthanized 6 h and 24 h post-mAb injection. Data are the mean values (tumor/organ) with SD (n =5).

| **Tumor/organ** | **6 h no trigger** | **6 h trigger 10** | **24 h no trigger** | **24 h trigger 10** |
|---|---|---|---|---|
| blood | 1.0 ± 0.4 | 2.3 ± 0.6 | 2.5 ± 0.7 | 6.6 ± 0.9 |
| heart | 3.8 ± 1.3 | 8.9 ± 2.3 | 9.0 ± 2.4 | 22.44 ± 3.0 |
| lung | 2.2 ± 0.7 | 4.0 ± 1.0 | 4.8 ± 1.8 | 10.8 ± 2.9 |
| liver | 3.8 ± 1.1 | 6.0 ± 1.9 | 9.3 ± 2.2 | 13.6 ± 3.4 |
| spleen | 4.9 ± 1.4 | 10.8 ± 3.6 | 11.4 ± 4.0 | 20.5 ± 4.7 |
| pancreas | 12.3 ± 5.4 | 21.2 ± 6.7 | 19.0 ± 6.4 | 36.6 ± 6.1 |
| kidney left | 3.1 ± 1.3 | 3.8 ± 1.2 | 5.4 ± 1.7 | 6.4 ± 1.2 |
| kidney right | 2.9 ± 1.3 | 3.6 ± 0.7 | 5.5 ± 1.2 | 6.4 ± 1.1 |
| muscle | 24.1 ± 10.0 | 34.7 ± 11.9 | 31.8 ± 11.4 | 67.9 ± 21.8 |
| bone | 9.9 ± 4.0 | 13.6 ± 4.7 | 12.8 ± 3.6 | 16.6 ± 3.5 |
| brain | 45.8 ± 19.3 | 85.9 ± 24.7 | 106.3 ± 29.2 | 218.8 ± 33.5 |
| fat | 5.1 ± 1.6 | 8.6 ± 3.2 | 7.7 ± 1.3 | 15.8 ± 4.5 |
| skin | 4.3 ± 2.2 | 7.4 ± 2.0 | 7.9 ± 2.9 | 15.3 ± 3.2 |

### *References*

1. Rossin R, Van Duijnhoven SMJ, Ten Hoeve W, et al. Triggered Drug Release from an Antibody-Drug Conjugate Using Fast ‘click-to-Release’ Chemistry in Mice. Bioconjugate Chemistry. 2016; 27: 1697–706.

2. Wang Q, Wang Y, Ding J, et al. A bioorthogonal system reveals antitumour immune function of pyroptosis. Nature. 2020; 579: 421–6.
